# Can Sexual Selection Cause Divergence in Mating System-Related Floral Traits?

**DOI:** 10.1101/513481

**Authors:** Åsa Lankinen, Maria Strandh

## Abstract

**Premise of the Research:** The wide diversity of floral traits seen among plants is shaped by neutral and selective evolutionary processes. In outcrossing species, sexual selection from competing pollen donors is expected to be important for shaping mating system-related traits but empirical evidence is scarce. In a previous evaluation of experimental evolution lines crossed with either one or two pollen donors (monogamous, M, or polyandrous, P, lines) at early floral stages in mixed-mating *Collinsia heterophylla* (Plantaginaceae), P showed enhanced pollen competitive ability and reduced maternal seed set compared to M, in accordance with sexually antagonistic evolution of pollen. Here, we asked whether the presence of sexual selection during pollen competition affect mating system-related floral traits in the same lines.

**Methodology:** We compared flowering start, timing of anther-stigma contact (as an indication of timing of self-pollination), timing of stigma receptivity and first seed set between M and P, and with a source line, S (starting material). The former three traits are later in outcrossers than in selfers of *Collinsia*. The latter trait was expected to be earlier in P than in M because of sexual selection for early seed siring of pollen.

**Pivotal Results:** Artificial polyandry for four generations resulted in later flowering start and later anther-stigma contact in P compared to M, and the latter trait was intermediate in S. Thus, P appeared more ‘outcrossing’ than M. Stigma receptivity did not differ between lines. First seed set was earlier in P than in M, as expected from sexual selection.

**Conclusions:** Our results from *C. heterophylla* experimental evolution lines suggest that a component of sexual selection during outcross pollination could enhance the patterns of floral divergence commonly found between outcrossers and selfers.

## Introduction

The wide floral diversity in angiosperms is considered largely to be caused by geographic variation in pollinator-mediated selection (Kay and Sargent 2009; Van Der Niet et al. 2014; Armbruster 2014). Because differences in mating system (e.g. outcrossing vs. self-fertilization, or their combination in mixed mating) is correlated with floral and developmental traits (Karron et al. 2012; Barrett 2013), selective forces influencing mating system also contribute to angiosperm floral diversity. Traits such as small flower size and reduced separation of male-female functions in space and time (herkogamy and dichogamy) are thought to directly favour selfing as reproductive assurance when pollinator visits are unpredictable (Lloyd 1979; Lloyd and Schoen 1992; Opedal 2018). These traits can also be connected to rapid maturation or reduced investment in cross-pollination in selfers (Snell and Aarssen 2005; Sicard and Lenhard 2011).

It has recently been argued that not only maternal outcrossing rate but also mate diversity and individual variation in mating success could contribute to mating system selection (Barrett and Harder 2017). An influence of sexual selection on mating system evolution could be important to consider for a more complete understanding of the selective forces affecting mating system, because it is expected that sexual selection should be relatively more important in outcrossing taxa than in selfing taxa (Mazer et al. 2010). For example, due to stronger sexual selection or parental conflicts in outcrossers than in selfers, previous studies have suggested that pollen competitive ability during pollen competition in the pistil (Mazer et al. 2018), pistil barriers to hybridization (Brandvain and Haig 2005) and male vs. female antagonistic influence on seed provisioning (Willi 2013) are increased in outcrossers compared to in selfers. While sexual selection can influence floral traits (Delph and Ashman 2006; Moore and Pannell 2011; Dai and Galloway 2013), the contribution of this selective force on floral trait divergence between outcrossers and selfers is usually not considered.

The mixed-mating herb *Collinsia heterophylla* belongs to a genus with extensive variation in mating system, from self-pollinating to mixed-mating species (Armbruster et al. 2002; Kalisz et al. 2012). Most outcrossing species have larger flowers, delayed selfing brought about by loss of herkogamy at late developmental stages and delayed stigma receptivity, separating timing of male and female reproductive functions (fig. 1a). These floral traits also appear to be associated with slower plant developmental rate and later flowering start (Elle et al. 2010). In *C. heterophylla*, variation in outcrossing rate is substantial and populations with higher outcrossing rates have later timing of stigma receptivity (Strandh et al. 2017). In line with the results at the genus level, delayed selfing measured as timing of anther-stigma contact was shown to be genetically correlated with timing of stigma receptivity, and tended to be genetically correlated with flowering start (Lankinen et al. 2017a).

**Fig. 1.**
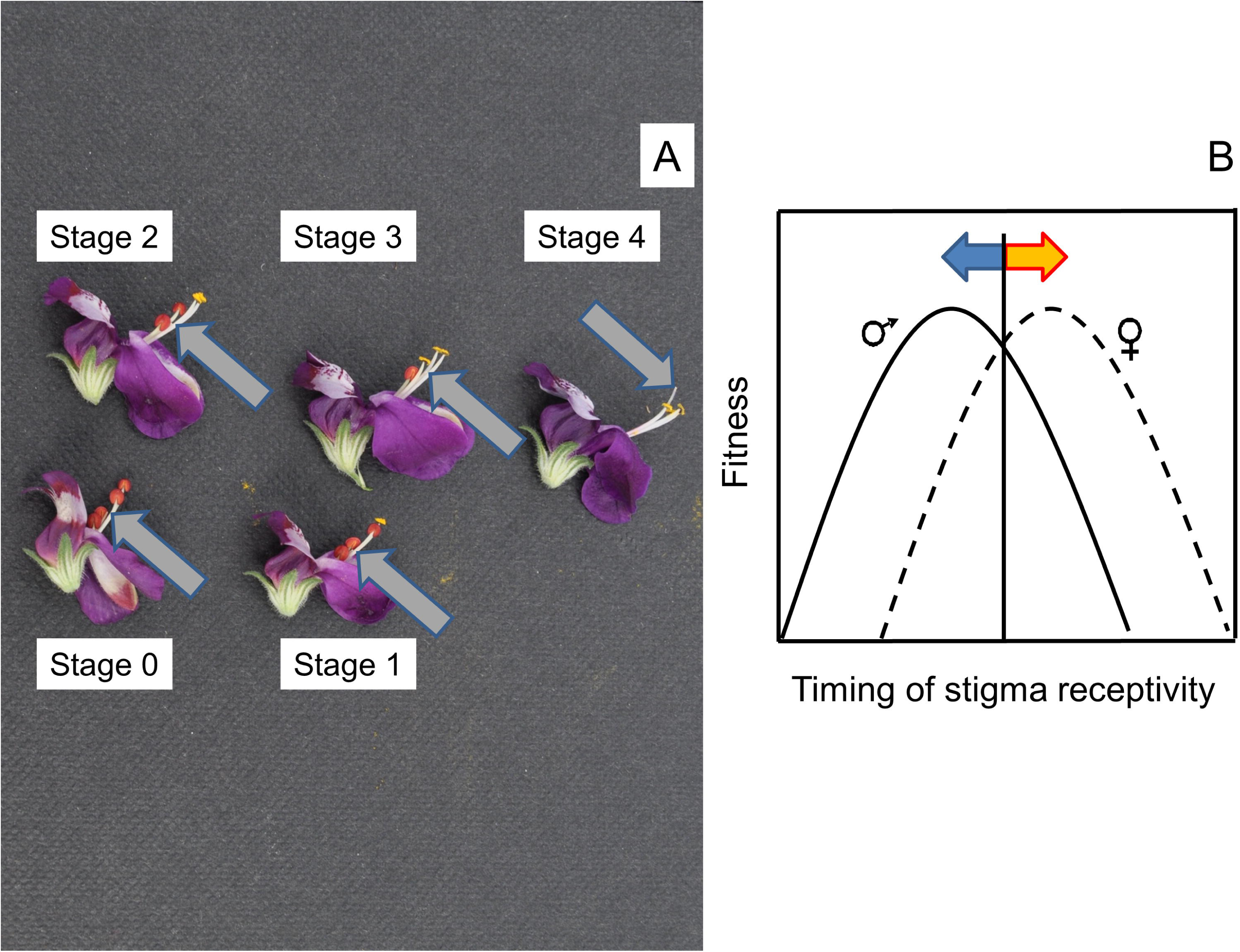
A) Floral developmental at stage 0-4 in *Collinsia heterophylla*. During floral development, the anthers dehisce and the pistil elongates. At later stages, stigma receptivity occurs and the pistil grows through its own pollen, allowing delayed selfing. Stage 0 = stage of flower opening, stage 1-4 = stage with number of dehisced anthers, respectively. Gray arrows point to the location of the stigma. B) Model of sexual conflict over timing of stigma receptivity, involving opposing selection pressures in male and female function due to their divergent evolutionary interests. Moving trait values of the male function closer to its fitness optimum (blue arrow) in *C. heterophylla* in an experimental evolution experiment, caused a direct fitness cost in the female function in terms of reduced seed set (Lankinen et al. 2017b). Photograph in A): Josefin A Madjidian.

The timing of stigma receptivity has been proposed to be affected by a sexual conflict in *C. heterophylla* (Lankinen and Kiboi 2007) (fig. 1b). A sexual conflict involves opposing selection pressures in males and females due to divergent evolutionary interests of the sexes (Parker 1979). When such conflicts occur between different loci in the two sexes (interlocus conflict), selection is expected to move trait values of one sex closer to its fitness optimum, causing a direct fitness cost in the other sex (Parker 1979; Arnqvist and Rowe 2005). Sexual selection for a trait value leading to increased reproductive success can generate sexual conflict (Kokko and Jennions 2014). The sexual conflict over timing of stigma receptivity in *C. heterophylla* involve i) pollen ability to sire seeds early to secure paternity when stigmas are partially receptive, and ii) a recipient cost of reduced early seed set (Lankinen and Kiboi 2007; Madjidian et al. 2012). We recently studied the evolutionary outcome of this conflict by producing experimental evolution lines by crossing recipients at early floral development with two pollen donors (polyandrous, P) or with one pollen donor (monogamous, M) (fig. 2a) for four generations (Lankinen et al. 2017b). Recipients always contributed with one offspring to the next generation, thus limiting selection on recipients compared to on pollen donors. We showed that P plants produced pollen with a higher proportion of successful crosses at early floral stages and faster tube-growth rate, and reduced seed set compared to M plants Lankinen et al. 2017b). These results are in accordance with enhanced sexual conflict and antagonistic evolution of P pollen in response to sexual selection (Arnqvist and Rowe 2005).

**Fig. 2.**
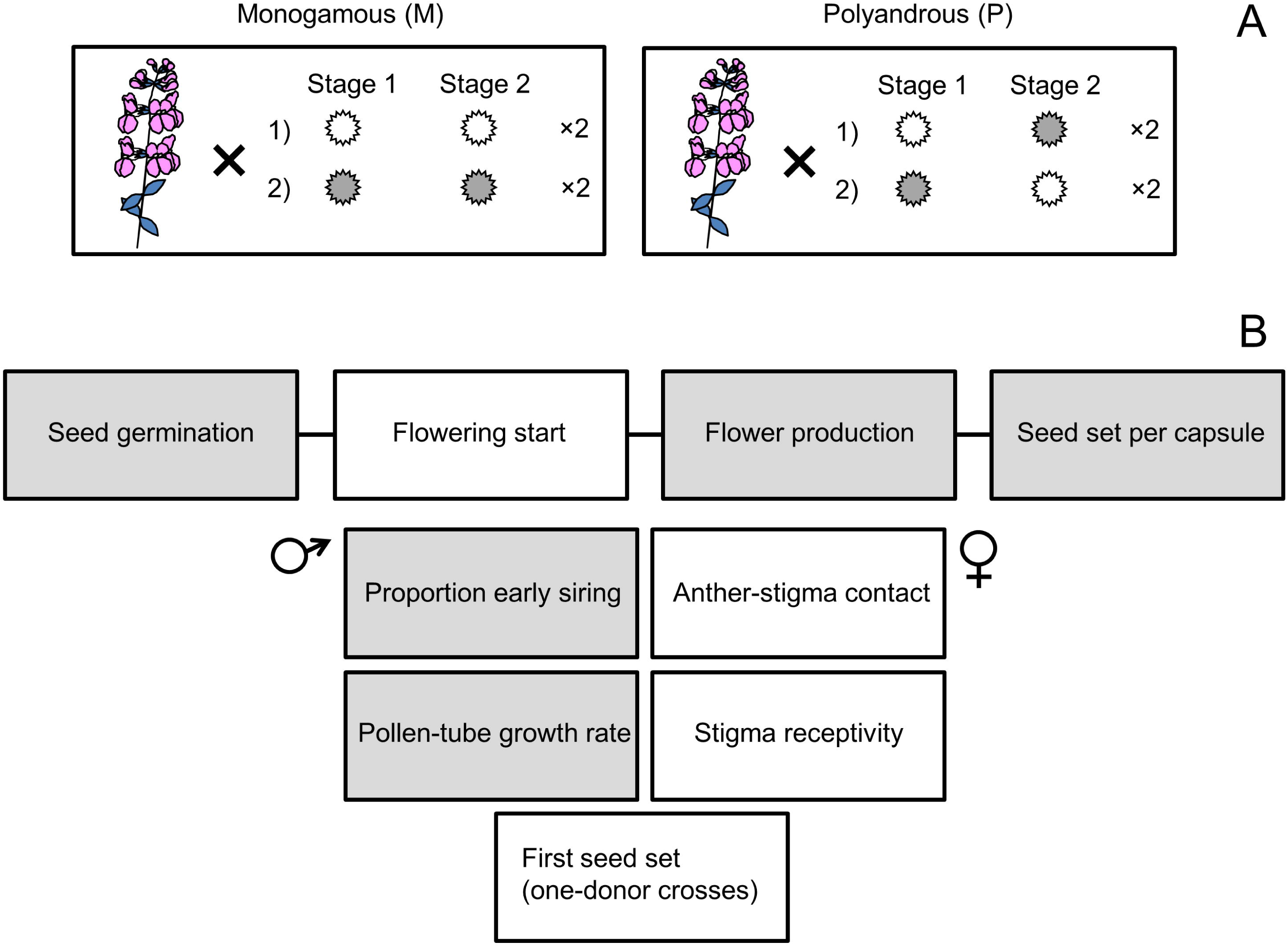
Overview of experimental design and results of experimental evolution in *Collinsia heterophylla*. A) Experimental evolution lines M (monogamous, one pollen donor) and P (polyandrous, two pollen donors) were produced by controlled hand-pollinations in the greenhouse for four generations starting from S (source) line. In both M and P, four flowers per recipient were repeatedly crossed at stage 1 and 2 (unreceptive or partially receptive pistils) involving two pollen donors (gray vs. white pollen grains). In M, each flower received pollen from the same donor while in P, each flower received flowers from two different donors. B) Overview of estimated traits related to different life stages and mating system in S, M and P lines. Pollen traits (proportion early siring and pollen-tube growth rate) represent the male reproductive function and pistil traits (anther-stigma contact and stigma receptivity) represent the female reproductive function. First seed set is the result of the combined influence of male-based and female-based influence on stigma receptivity and seed set. Results for traits in white boxes = reported in the current study, gray boxes = reported in Lankinen et al. (2017b).

In the current study, we continued analysing the experimental evolution lines in *C. heterophylla* (Lankinen et al. 2017b), asking whether the presence of sexual selection during pollen competition also affected mating system-related floral traits (fig. 2b). Because of the importance of timing of stigma receptivity in relation to both mating system and sexual conflict in *C. heteropylla*, we hypothesized that sexual selection on early pollen competitive ability could affect this trait as well as two other correlated floral traits, flowering start and timing of anther-stigma contact. We hypothesized that the P line would show later flowering start, later anther-stigma contact, and later stigma receptivity, but earlier timing of first seed set, i.e. stigma receptivity influenced by both pollen and pistil (Table 1). The prediction for later stigma receptivity is based on our previous finding of a negative correlation between male and female influence on first seed set (Hersh et al. 2015), suggesting later stigma receptivity when selecting for earlier pollen influence on stigma receptivity, given that this association is genetically determined. We also compared M and P to a source line, S, representing starting material outcrossed at late floral stages for one generation Lankinen et al. 2017b), i.e. S was not selected for early performance of pollen and pistil traits. A comparison with S could give an indication of the direction of evolutionary change of M and P from the source.

**Table 1.**
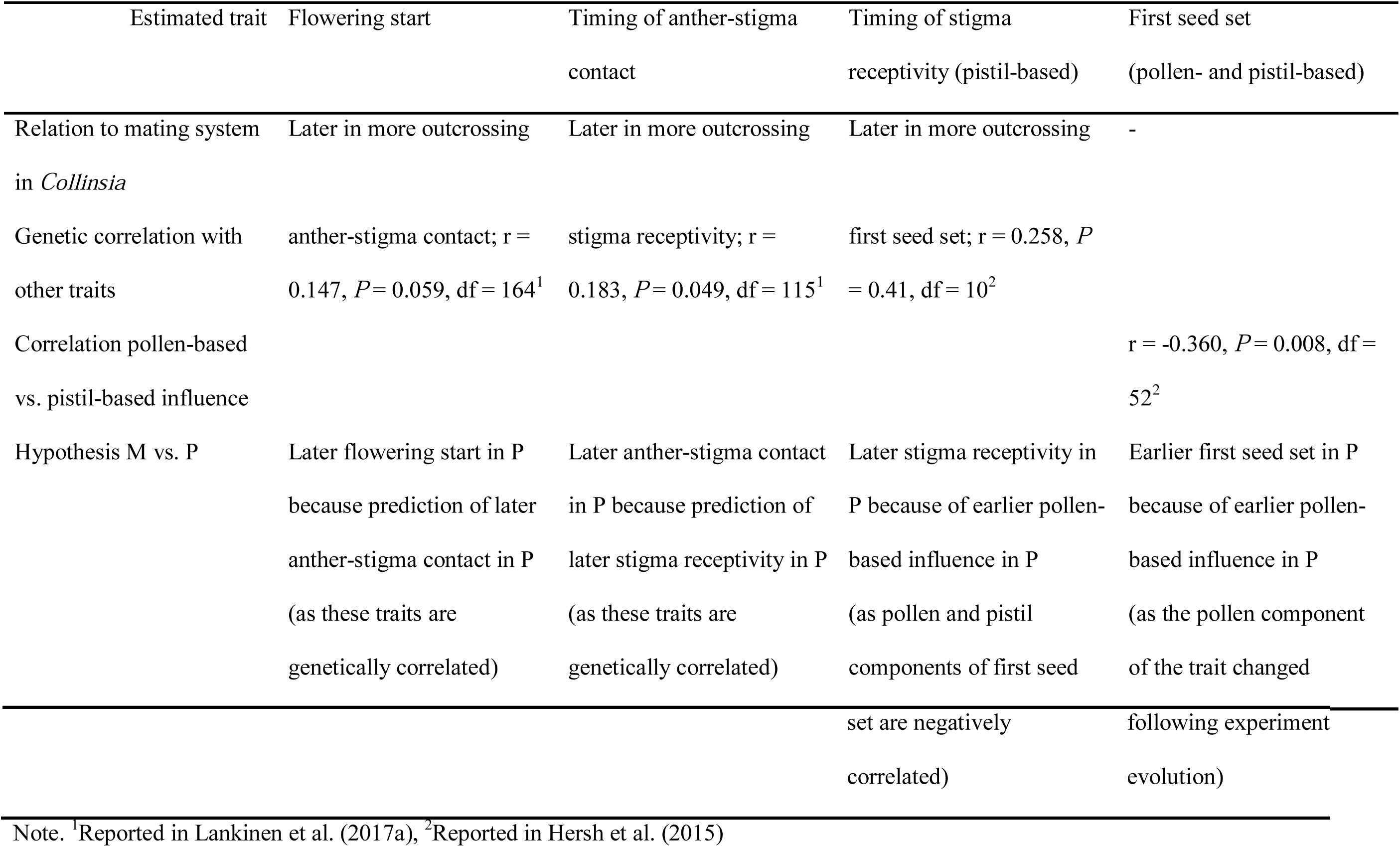
Previous results and hypotheses tested in the present study regarding floral traits estimated among M (monogamous), P (polyandrous) and S (source) experimental lines in *Collinsia heterophylla*

## Materials and methods

### Study species and experimental evolution lines

*Collinsia heterophylla* Buist (Plantaginaceae) is a hermaphroditic, self-compatible winter-annual herb native to California (Newson 1929; Neese 1993). The species is pollinated by long-tongued, nectar-feeding bees (Armbruster et al. 2002). Flowers are zygomorphic with five-lobed corollas forming an upper and a lower lip arranged in whorls on spikes. They contain four epipetalous stamens and one single-style pistil that develop into seed capsules containing up to 20 seeds (Armbruster et al. 2002; Madjidian and Lankinen 2009). During floral development, the four anthers mature and dehisce at a rate of approximately one per day during four consecutive days (fig. 1a). The stigma becomes receptive around day 2-3 after flower opening and the style elongates, placing the stigma in contact with the mature anthers providing an opportunity for delayed selfing at about the same time as stigma receptivity (Armbruster et al. 2002; Madjidian and Lankinen 2009) (fig. 1a).

Plants used in this experiment originated from seeds collected by maternal family (*N* = 200) in a large natural population in Mariposa county (situated at N 37.50196; W 120.12360) in 2008. As described in Lankinen et al. (2017b), we used this material to create an outcrossed source (S, *N* = 177 maternal families) population in 2010 for our experimental evolution study in 2010-2013. We performed controlled hand-pollinations at floral developmental stage four, representing day four after flower opening, i.e. when stigmas were fully receptive. Flowers were emasculated at flower opening (= stage zero). The traits investigated in Lankinen et al. (2017b) and anther-stigma contact, stigma receptivity and first seed set estimated in the current study (fig. 2b) were similar to previous greenhouse studies using plant material from the same population (Madjidian and Lankinen 2009; Hersh et al. 2015), despite storage of seeds.

From S we created two evolved lines; i) monogamous line (M, *N* = 135 maternal families) crossed with one pollen donor and ii) polyandrous line (P, *N* = 142 maternal families) crossed with two pollen donors (fig. 2a) by conducting hand-pollinations on emasculated flowers for four generations in 2010-2012 (Lankinen et al. 2017b). The evolved lines were unreplicated. We sampled a large number of genotypes within each treatment to capture a high degree of the natural genetic variation. This will minimize the impact of genetic drift in experimental evolution (Fuller et al. 2005). To impose sexual selection on early siring success of pollen in partially receptive pistils, we conducted crosses twice per flower at early floral stages 1 and 2, i.e. day 1 and 2 after flower opening (fig. 2a). We hand-pollinated four flowers per recipient plant involving two different pollen donors (see a more detailed description in Lankinen et al. (2017b)). One of the seeds generated per plant gave rise to the next generation, thus reducing selection on the female reproductive function compared to selection on the male function. All experimental plants were raised from cold-stratified seeds and grown in an insect-free greenhouse.

### Estimates of floral traits among experimental lines

We assessed four floral traits of the S line as well as of the two evolved lines M and P (fig. 2b). We recorded flowering start as number of days since the first plant, independent of line, started flowering (*N* families = 21 in S, 25 in M, 29 in P, *N* = 3-8 plants per family). Timing of anther-stigma contact, as an indication of timing of self-pollination, was assessed by noting the floral developmental stage (= number of dehisced anthers) when the stigma was in contact with the open anthers (*N* families = 20-22 per line, *N* = 2 plants per family, *N* = 2 flowers per plant and floral stage 1-4). Timing of stigma receptivity was determined in a droplet of 3% hydrogen peroxide (Kearns and Inouye 1993) in emasculated flowers at stage 1-4 (=day after flower opening) (*N* families = 12-17 per line, *N* = 1 plants per family, *N* = 1-2 flowers per plant and floral stage). Vigorous bubbling on the stigmatic surface (unharmed and pollen free tissue) suggests activity of stigmatic peroxidase, which has been shown to correlate with presence of pollen tubes in the pistil following hand-pollination in this species (Lankinen et al. 2007).

Timing of first seed set was calculated from one-donor hand-pollinations performed in emasculated flowers at day 1-4 after flower opening (*N* = 9 recipients and 6 pollen donors per line, *N* = 16 crosses per recipient involving 2 donors, and 2 flowers per stage and donor) (Lankinen et al. 2017b). Four h after the crosses the stigma and upper part of the style were removed to ensure that seed formation only occurred in flowers with stigmas receptive at the time of the cross.

We analyzed differences in the measured traits i) among S, M and P, or ii) M and P using ANOVAs (type III sums of squares) in SPSS (SPSS 2016). Because our main focus was to investigate M and P divergence, the latter analysis was performed when no significant differences were detected among the three lines, to evaluate differences between M and P potentially masked by inclusion of S. When more than one plant was estimated per family, we used a nested model including line (fixed) and family (random) nested within line. In other cases, we only included line. Differences among lines were determined by Tukey tests. Timing of first seed set was square-root transformed. Models were evaluated for normality and homogeneity of the residuals.

## Results

Flowering start and timing of anther-stigma contact showed divergence between M and P following four generations of experimental evolution (Table 2, M-P: flowering start, *P* = 0.034; anther-stigma contact, *P* = 0.001). In accordance with our hypotheses, both traits were later in P than in M (fig. 3a,b). Our source S had earlier flowering start than the evolved lines (S-M and S-P: *P* < 0.001, fig. 3a), but no difference was found for timing of anther-stigma contact (*P* > 0.13, fig. 3b).

**Fig. 3.**
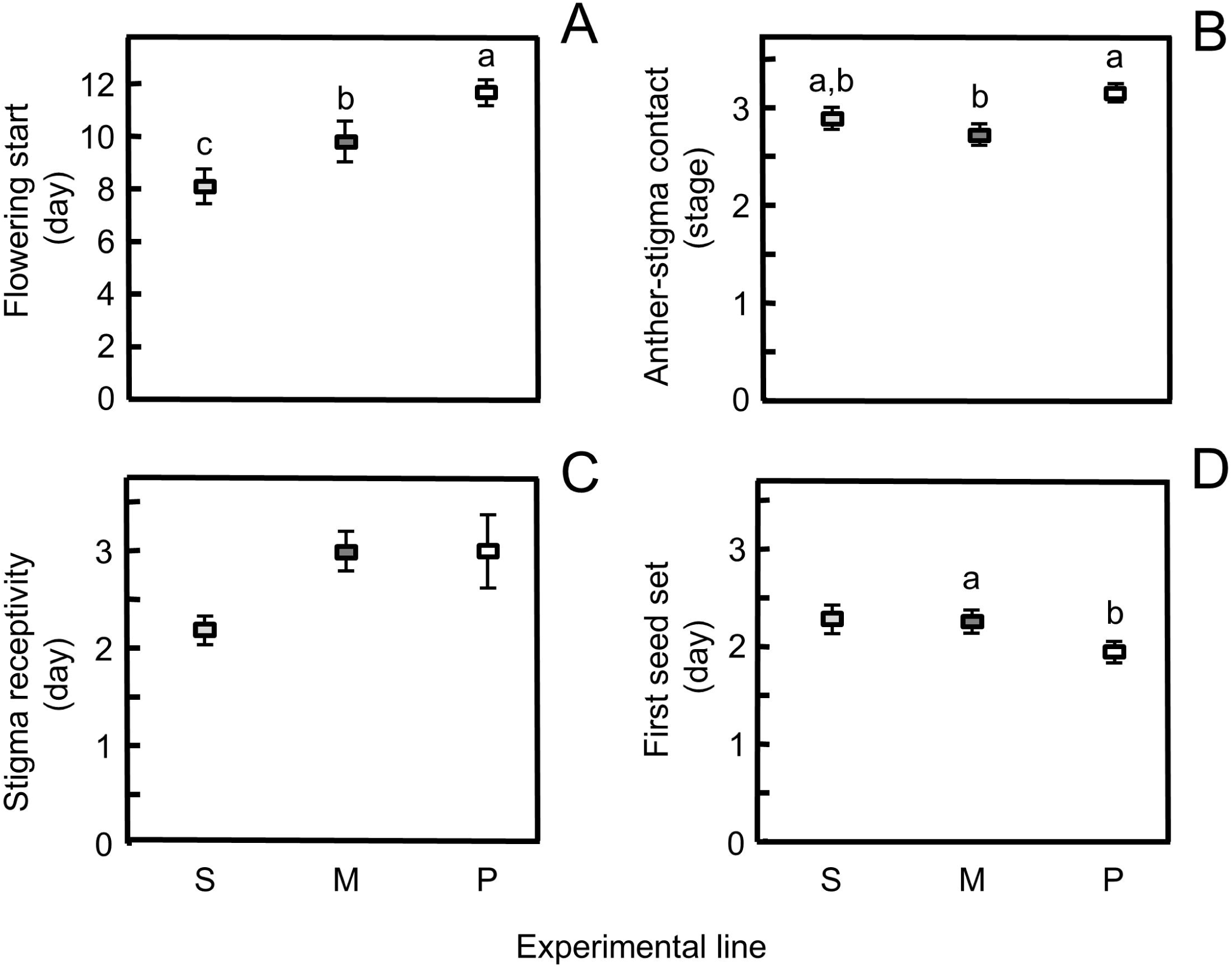
Floral traits related to mating system in *Collinsia heterophylla* among M (monogamous), P (polyandrous) and S (source) experimental lines averaged over recipients and when necessary over pollen donors. A) Flowering start estimated as day from flowering of the first plant independent of line. B) Anther-stigma contact estimated as floral developmental stage (= number of dehisced anthers) when the stigma grows into the dehisced anthers. C) Stigma receptivity estimated as day after flower opening when peroxidase activity occurs. D) First seed set estimated as day after flower opening when seeds are formed following controlled hand-pollinations and subsequent pistil removal at day 1-4 after flower opening. Error bars indicate ±1 SE. Different letters denote significant difference for tests performed among the three lines (A-B) and between M and P (D).

**Table 2.** Analyses of variance of floral traits related to mating system among M (monogamous), P (polyandrous) and S (source) experimental lines in *Collinsia heterophylla*

|  | Flowering start |  |  | Anther-stigma contact |  |  | Stigma receptivity |  |  | First seed set |  |  |
| --- | --- | --- | --- | --- | --- | --- | --- | --- | --- | --- | --- | --- |
|  | (day) |  |  | (stage) |  |  | (day) |  |  | (day) |  |  |
| Source of variation | df | *F* | *P* | df | *F* | *P* | df | *F* | *P* | df | *F* | *P* |
| Line | 2,74.9 | 6.93 | 0.002 | 2,61.5 | 4.31 | 0.018 | 2,39 | 2.66 | 0.082 | 2,24.1 | 2.05 | 0.15 |
| Recipient nested within line | 72,354 | 6.33 | <0.001 | 61,63 | 1.628 | 0.028 |  |  |  | 24,80 | 1.01 | 0.47 |

Contrary to our hypothesis, timing of stigma receptivity did not differ among all three lines (Table 2, fig. 3c) or between M and P (*F*_1,28_ = 0.026, *P* = 0.87). Variability was higher in P than in M (*F*-test; *P* = 0.037, *N* = 30) and in S (*P* = 0.002, *N* = 26), which was an unexpected result.

Day of first seed set following one-donor pollinations, showed, as predicted, earlier formation of seeds in P than in M (Line: *F*_1,16_ = 4.90, *P* = 0.042, Recipient nested within line: *F*_16,54_ = 0.691, *P* = 0.79, fig. 3d). There was no significant difference among all three lines (Table 2).

## Discussion

While sexual selection has been studied in plants for several decades, this mode of selection is still not a well-integrated concept in plant evolution theory, including the theory on mating system evolution (Lankinen and Karlsson Green 2015). In the current study we investigated a potential link between sexual selection and floral traits related to the mating system in *C. heterophylla*, a mixed mating species belonging to a genus with extensive variation in mating system and associated floral traits (Armbruster et al. 2002; Kalisz et al. 2012). To study how the presence of sexual selection impacts evolution of floral traits, we compared monogamous (M) and polyandrous (P) experimental evolution lines, both outcrossed at early floral stages (Lankinen et al. 2017b). A previous analysis of M and P showed that the presence of sexual selection in P led to higher levels of sexual conflict with increased pollen competitive ability and reduced seed set. With the help of other previous studies in *C. heterophylla* on genetic and phenotypic correlations between floral traits (Hersh et al. 2015; Lankinen et al. 2017a) we made specific predictions regarding the estimated traits flowering start, timing of anther-stigma receptivity, stigma receptivity (all three later in P vs M, Table 1) and first seed set (earlier in P vs M, Table 1).

In *C. heterophylla*, flowering start and timing of anther-stigma contact, the latter as an indication of timing of self pollination, showed divergence between M and P. Both traits were later in P than in M, which was in line with expectations from genetic correlations between these two traits and between timing of anther-stigma contact and stigma receptivity (Lankinen et al. 2017a, Table 1). Thus, P appeared more ‘outcrossing’ than M (Elle et al. 2010; Kalisz et al. 2012; Strandh et al. 2017). The source line (S) had earlier flowering start than both M and P. This result may imply evolution of later flowering start in both M and P, which were produced at early floral stages compared to S. We cannot, however, exclude that the earlier flowering start in S could be related to the previously found reduction in seed germination rate and number of flowers in S (Lankinen et al. 2017b), which may have resulted from longer storage of S seeds than seeds from M and P. Anther-stigma contact in S was intermediate between M and P. For this trait S was similar to that found in other greenhouse studies using plant material from the same population (Madjidian and Lankinen 2009). This suggests that divergence in anther-stigma contact was caused by P becoming later and M becoming earlier than S. We can hypothezise that not only the presence of sexual selection but also the absence of sexual selection during outcrossing can impact divergence of pollen and floral traits. Interestingly, in *Clarkia xantiana* strong pollen limitation resulted in disruptive selection through female and male fitness, as reduced herkogamy and protandry increased female fitness while both large and small petal area increased male fitness (Briscoe Runquist et al. 2017). It would be highly informative with more studies on the influence of sexual selection on mating system-related traits also in other study systems.

Timing of stigma receptivity did not differ significantly between M and P. This result was contrary to expectation (Table 1). First seed set, a proxy for stigma receptivity influenced by both pistil and pollen, was earlier in P than in M, as predicted from the previous analyses of M and P (Lankinen et al. 2017b) suggesting that P pollen was more successful at siring seeds in partially receptive pistils. This result, in combination with a detected negative relationship between male and female influence on first seed set within individual plants (Hersh et al. 2015), indicated that we could expect later stigma receptivity in P. While the mean of stigma receptivity did not differ between M and P, variability of this trait was higher in P compared to in both M and S. This result was surprising. One possible explanation is that there was disruptive selection acting on this trait in P, favoring either early or late stigma receptivity. In *Drosophila melanogaster*, experimental evolution, involving disruptive selection by alternating up and down selection, increased phenotypic variation in wing shape, while fluctuating and stabilizing selection instead decreased the variation (Pélabon et al. 2010). We do not have support for disruptive selection on stigma receptivity. We could, however, hypothesize contrasting effects of direct influence of pollen (early pollen-based influence will lead to earlier receptivity, Lankinen et al. 2017b) and indirect genetic covariance (negative correlation between pollen- and pistil-based influence on stigma receptivity, Hersh et al. 2015). Interestingly, several previous studies in *C. heterophylla* found that stigma receptivity is more variable than anther-stigma contact (Lankinen et al. 2007; Madjidian and Lankinen 2009; Hersh et al. 2015).

Despite that timing of stigma receptivity was not significantly later in P, we surmise that the later response in both flowering start and timing of anther-stigma contact in P was a consequence of genetic correlations among traits (Lankinen et al. 2017a). While flowering start was not significantly genetically correlated with the other two traits (Lankinen et al. 2017a), it is possible that these three traits are genetically linked with plant developmental rate. Other studies indicate that selection for rapid development can be correlated with rapid flower maturation (Mazer et al. 2004; Snell and Aarssen 2005; Elle et al. 2010). However, a recent study suggested low levels of genetic covariances between herkogamy and other floral traits across 17 species representing 10 families (Opedal et al. 2017). Evolvability of herkogamy was estimated to be 9.07%, which was an order of magnitude greater than evolvabilities of the male and female organs that are components of herkogamy, and of flower size. Because these results suggest that herkogamy has a high potential to respond to natural selection, it is possible that this trait is only weakly influenced by a component of sexual selection in other study systems.

A weakness of our study is the lack of replication of the experimental evolution lines. This implies that we cannot fully exclude genetic drift as a cause of the results, despite following the recommendation of a large number of individuals within each line to reduce the influence of drift (Fuller et al. 2005). However, several additional experiments suggest the presence of a sexual conflict over timing of stigma receptivity in *C. heterophylla* (Lankinen and Kiboi 2007; Madjidian and Lankinen 2009; Madjidian et al. 2012; Hersh et al. 2015), indicating that the reported differences between M and P in pollen competitive ability and seed set is caused by sexual selection rather than genetic drift (Lankinen et al. 2017b). Moreover, the link between floral traits is well documented both in *C. heterophylla* (Lankinen et al. 2017a) and among species in *Collinsia* (Armbruster et al. 2002; Kalisz et al. 2012), further supporting a scenario of selection rather than drift.

In conclusion, the influence of sexual selection in *C. heterophylla* experimental evolution lines resulted in divergence of mating system-related floral traits, at least for timing of anther-stigma contact, which is related to timing of self pollination and herkogamy. P appeared more ‘outcrossing’ than M. Thus, adding a component of sexual selection during outcross pollination could enhance the patterns of floral divergence regularly seen between selfers and outcrossers (Karron *et al.*, 2012; Barrett, 2013). It would be of great interest to learn if sexual selection could impact divergence in herkogamy, or other mating system-related floral traits, also in other study systems. Such knowledge could lead to a better understanding of how multiple factors influence evolution of plant mating system and floral divergence. In line with Barrett and Harder (2017), we suggest that future studies should consider sexual selection and mate diversity in relation to plant mating system selection and divergence, and the possibility that some of the floral trait divergence we see today between outcrossers and selfers is in fact a result of sexual selection.

## Acknowledgements

We thank S Hydbom for assistance in the greenhouse. This work was supported by the Carl Trygger Foundation for Scientific Research; the Crafoord Foundation and the Swedish Research Council (to ÅL).

